# ConfuseNN: Interpreting convolutional neural network inferences in population genomics with data shuffling

**DOI:** 10.1101/2025.03.24.644668

**Authors:** Linh N. Tran, David Castellano, Ryan N. Gutenkunst

## Abstract

Convolutional neural networks (CNNs) have become powerful tools for population genomic inference, yet understanding which genomic features drive their performance remains challenging. We introduce ConfuseNN, a method that systematically shuffles input haplotype matrices to disrupt specific population genetic features and evaluate their contribution to CNN performance. By sequentially removing signals from linkage disequilibrium, allele frequency, and other population genetic patterns in test data, we evaluate how each feature contributes to CNN performance. We applied ConfuseNN to three published CNNs for demographic history and selection inference, confirming the importance of specific data features and identifying limitations of network architecture and of simulated training and testing data design. ConfuseNN provides an accessible biologically motivated framework for interpreting CNN behavior across different tasks in population genetics, helping bridge the gap between powerful deep learning approaches and traditional population genetic theory.

## INTRODUCTION

In recent years, machine learning methods have seen tremendous growth in population genetics (see Schrider & Kern (2018); Huang et al. (2024) for recent reviews). Although classic population genetic inference approaches are based on summary statistics, the rapid advancement of computational capabilities coupled with increasingly sophisticated simulators has primed the field for a new paradigm, using supervised machine learning models trained on simulated data. The most popular among these algorithms thus far are convolutional neural networks (CNNs), which have been applied to a wide variety of population genetic inference tasks: inferring recombination rates and recombination hot spots (Chan et al. 2018; Flagel et al. 2019), demographic history (Flagel et al. 2019; Sanchez et al. 2021), hybrid speciation and admixture (Blischak et al. 2021), geographic dispersal (Smith et al. 2023), and detecting signature of introgression (Flagel et al. 2019; Gower et al. 2021) and natural selection (Flagel et al. 2019; Torada et al. 2019; Qin et al. 2022).

The exploration of CNN-based approaches in a wide range of tasks has yielded promising results: suites of trained CNNs with great reported performance on simulated data. The field is thus positioned for further investigation into the inner workings of these models to derive insights connecting performance to underlying population genetic processes and principles. Within genomic research, explainable AI (xAI) has also garnered great interest. Novakovsky et al. (2023) provides an in-depth review of the state-of-the-art approaches in post-hoc interpretation methods, which focus on the analysis of model performance and quantification of input feature importance after model training. In population genetics, while a few studies (Cecil & Sugden 2023; Riley et al. 2024; Hahn & Mishra 2025) have begun to open the “black box” and explore how these trained models operate, many questions remain. For example, what data features are being exploited by a CNN during training? Are there novel features in the data not captured by classic summary statistics that contribute to the accuracy? Does a CNN trained for a certain inference task become more sensitive to a certain data feature than a CNN trained for a different inference task?

To address these questions, we developed a new method for interpreting CNN performance in population genetics by data shuffling. Our method is straightforward to implement, because it does not require the complex network manipulations often employed in the field of xAI. Rather, we focus on specific manipulation of the input genomic data, disrupting features captured by common summary statistics in population genetics to gain insights into the performance of CNNs. We applied our method to three published CNNs trained for various population genetic inference tasks: Riley et al. (2024) (disc-pg-gan) and Torada et al. (2019) (ImaGene) for detecting signatures of positive selection, and Flagel et al. (2019) for inferring population demographic history. We show that our method reveals which population genetic data features are important for these CNNs to achieve their inference accuracy.

## RESULTS & DISCUSSION

### The genomic image pixel shuffling method

In a typical CNN used for population genetic inference, the input genetic variation data is represented as a 2D image with rows as haplotypes and segregating sites as columns, ordered from left to right according to the genomic positions of the segregating sites (Fig. 1A). Therefore, the way the pixels are organized in this image is intrinsically connected to classic summary statistics used to describe patterns of genetic variations. We used this insight to design a set of shuffling operations on these pixels, in which each operation specifically disrupts a feature of the data connected to a known summary statistic. We applied these operations to the test data (not the training data) and evaluated any decline in performance to assess which summary statistics matter most to the CNN.

**Figure 1:**
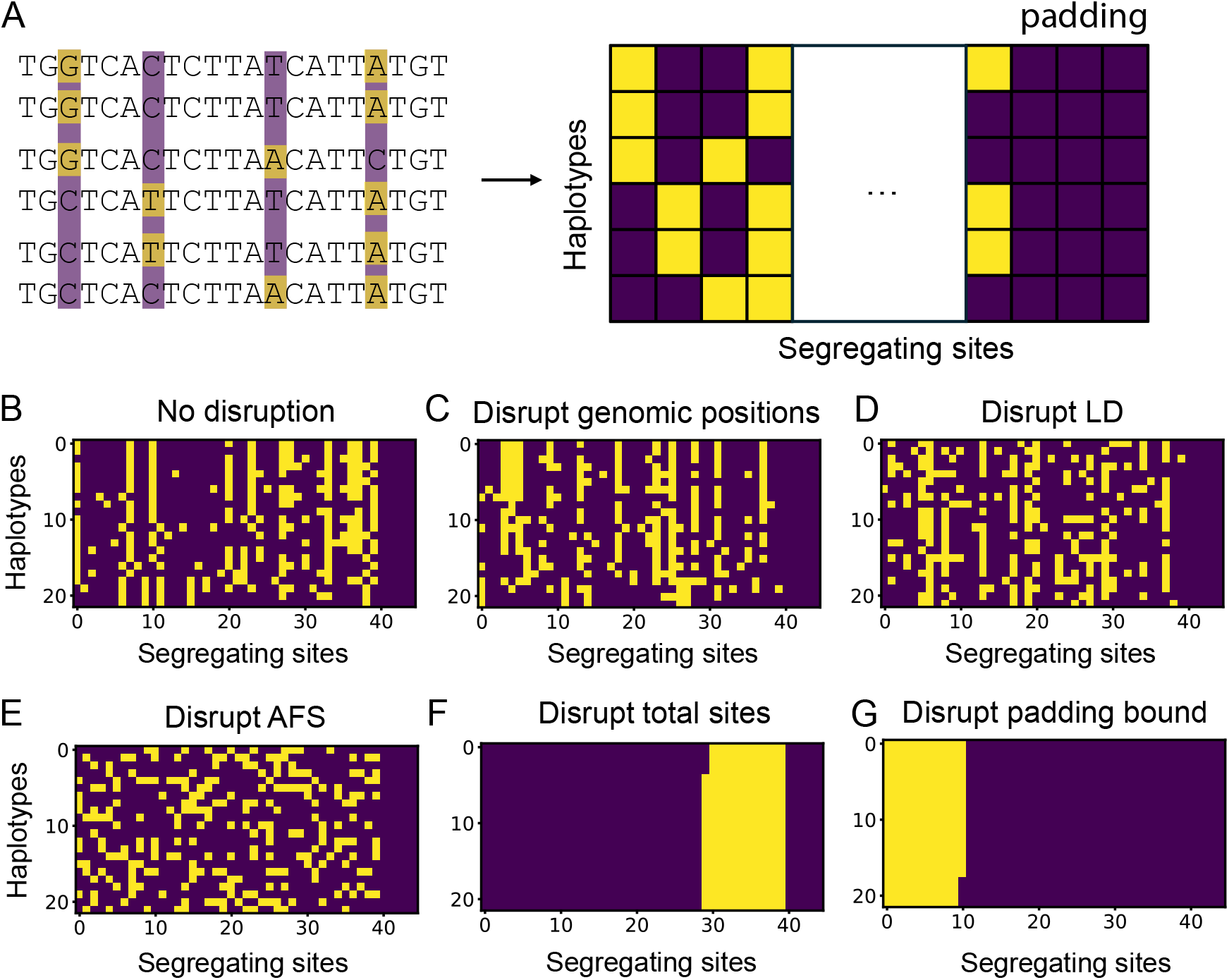
Illustration of the matrix representation of genetic variation data and the transformations of data shuffling. (A) From DNA sequence alignment to CNN input data matrix. In this example, six DNA sequences (haplotypes) are aligned and the biallelic segregating sites are highlighted by two different colors (left). The right-most all-purple columns represent the zero-padded portion of the data. (B-G) The data before and after the shuffling operations and the targeted features disrupted. (C) Randomly shuffling all columns of segregating sites in (B). (D) Randomly shuffling all entries within each column after shuffling columns as in (C). (E) Randomly shuffling all entries in (B). (F) Grouping entries of the same type into the same region, with the zero-padding maintained. (G) Same as (F) but entries are grouped in a way that disrupts the original zero-padding boundary.

The first and simplest operation is the random shuffling of full columns of segregating sites. This will alter the genomic positions of sites compared to the original data altering only short-scale linkage disequilibrium (LD)(Fig. 1B&C). Next, on top of the column shuffle operation, we can independently shuffle pixels within each column. These two operations together destroy correlations between neighboring sites and disrupt long-scale LD summary statistics (Fig. 1D). Another commonly used statistic in population genetics is the allele frequency spectrum (AFS), which summarizes the frequencies of genetic variants within the sample. To produce data with the AFS disrupted, we randomly shuffle all pixels (Fig. 1E). Finally, the most drastic disruption to the data groups all similar-color pixels into blocks. This operation disrupts the pattern of total diversity in addition to all aforementioned features (Fig. 1F&G). In addition to the haplotype matrix representation, some CNNs also require a vector of genomic positions or distances between the variants as an input. Our shuffling method does not make any changes to this vector.

### Application to published CNNs for population genetic inference

We applied our data shuffling procedure to three previously published CNNs designed to solve different inference problems in population genetics. Our general strategy was to regenerate the data used in the original study then apply the shuffles only to the test data. Data generation, preprocessing, and CNN training procedures were kept as similar to the original studies as possible.

Among the wide variety of population genetic inference tasks, detecting signatures of positive selection is one of the most popular. We investigated two CNNs designed for this task: Torada et al. (2019)’s Imagene and Riley et al. (2024)’s disc-pg-gan. Imagene was developed via classical supervised learning. By contrast, disc-pg-gan builds off a generative adversarial network framework for demographic history inference, in which the discriminator network was a CNN (Wang et al. 2021). That CNN was then fine-tuned with additional training on data simulated with selection to yield the disc-pg-gan network (Riley et al. 2024). Both CNNs were designed to classify whether the input image data contain a signal of adaptation, with the results presented here showcasing the performance of disc-pg-gan in binary classification (neutral or selection), and Imagene in multiclass classification (neutral, moderate selection, strong selection).

We applied the fine-tuned disc-pg-gan discriminator CNN to simulated test data, both similar to the original test and shuffled. As shown in Fig. 2A, disc-pg-gan’s CNN performance across the different sets of shuffled test data remains similar. The most drastic disruption of total sites only resulted in a slight reduction in performance. This suggests that the simulated test data may not have been challenging enough for fine-tuning the network. Specifically, the chosen test parameter regime generated data that were so different between the positive selection class and the neutral class that only learning total diversity (SNPs) from the data was enough for the network to achieve high accuracy. This finding is consistent with Riley et al. (2024)’s Fig. 3B showing that the pairwise heterozygosity statistic *p* has the highest correlation with the discriminator hidden units out of all the tested summary statistics. This result showcases how our shuffling method can detect problems with the simulation regime and provide direction to develop more appropriate test data.

**Figure 2:**
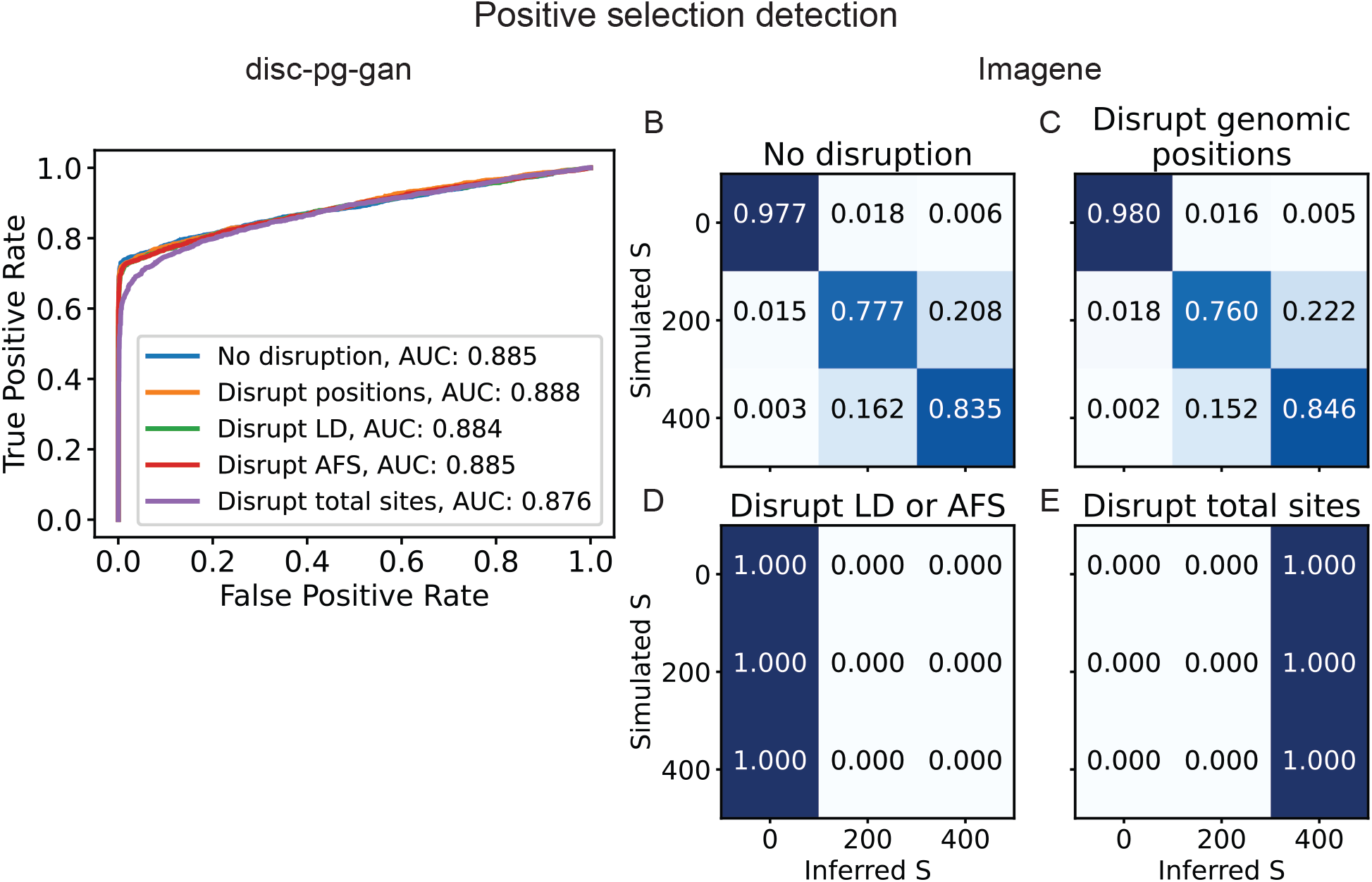
Performance of positive selection CNNs on genomic image data shuffled to remove known features. (A) Receiver operating characteristic (ROC) curve showing the performance of discpg-gan (Riley et al. 2024) inferring whether a genomic region is neutral or under positive selection. (B-E) Confusion matrices showing the performance of Imagene (Torada et al. 2019) inferring whether a genomic region is neutral (*S* = 0), under weak-to-moderate positive selection (*S* = 200), or under strong selection (*S* = 400).

**Figure 3:**
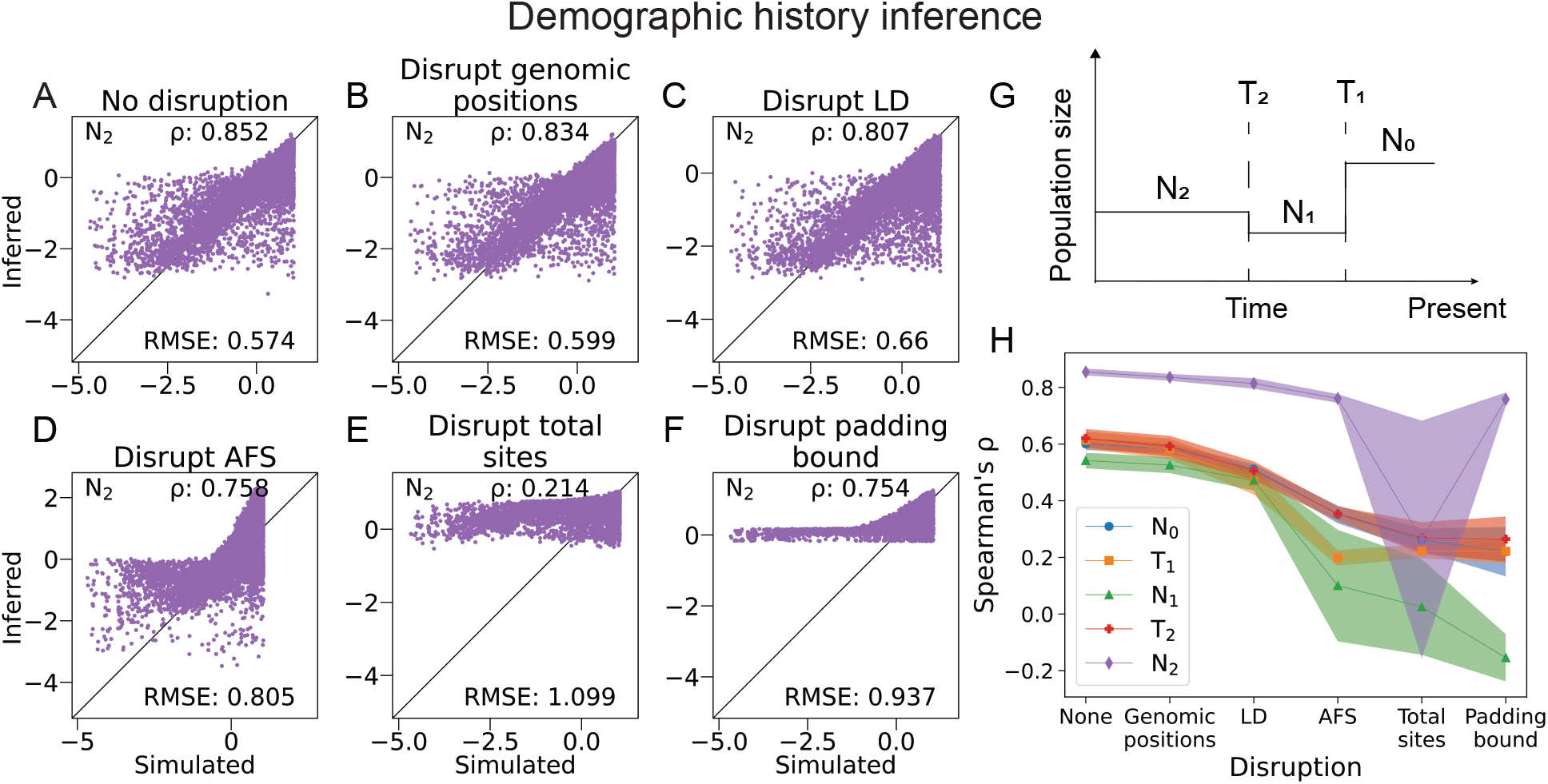
Performance of a CNN by Flagel et al. (2019) trained to infer parameters of a three-epoch demographic history model. This is a regression task for five demographic history model parameters: population sizes *N*_0_, *N*_1_, *N*_2_, and times of an instantaneous size change *T*_1_ and *T*_2_. (A-D) Scatter plots showing the correlation between simulated and inferred values using the trained CNN for the ancestral population size *N*_2_. Each dot represents a simulated data set, and Spearman’s *r* and RMSE scores summarize performance. Results for the other parameters are in Fig. S1. (G) Illustration of demographic history model inferred. (H) Patterns of performance degradation across shuffles are consistent among 10 independently trained instances of the CCN. Shaded regions indicate 95% confidence intervals based on the 10 instances.

For the multiclass classification task by Imagene, performance showed a clear breakdown when LD signals were disrupted (Fig. 2B-D). Further disruption at the AFS signal level did not change the pattern of failure. Interestingly, at the most drastic disruption level of total sites, the bias changed toward classifying all input data as experiencing strong selection instead of neutrality (Fig. 2D-E). This is consistent with intuition that this shuffling results in patterns in the data that resemble strong selection signals in which there is a dramatic reduction of haplotype diversity. Furthermore, disrupting linkage features greatly affected Imagene’s performance, indicating that this CNN likely relies on linkage features from the data to make its inference. A recent study (Cecil & Sugden 2023) further examining and breaking down Imagene found that performance similar to ImaGene can be achieved using just Garud’s H1 statistics (Garud et al. 2015), which is a LD statistic.

To further demonstrate the applicability of our shuffling method, we also applied it to a CNN developed for demographic history inference. Flagel et al. (2019) developed a CNN to infer five parameters for a three-epoch demographic history model (Fig. 3G). Here we tested the CNN architecture that the authors found to have the best accuracy performance among the many models that they built and tested.

We found that the perturbation of neither genomic positions nor LD had an effect on the performance of the CNN (Fig. 3A-C). In contrast, the demographic inference CNN performed significantly worse when the AFS signal in the data was disrupted, indicating the high impact of this feature on CNN learning for this task (Fig. 3D). Since this CNN includes zero-padding as a data preprocessing step, the position of the distinct allele block with respect to the padding dictates whether the total segregating sites boundary is preserved in the genomic data image (Fig. 1F-G). The demographic inference CNN performed much worse when this boundary was altered compared to when it was preserved while the other features were all disrupted (Fig. 3E&F), demonstrating the importance of this boundary to the CNN. We repeated the shuffle test for ten independently trained instances of this CNN architecture and found that the pattern of performance degradation is consistent across the instances (Fig. 3H).

Among the many CNN hyperparameters that Flagel et al. (2019) tried for the this inference task, the best performing convolutional kernel size was 2, meaning the convolutional filters only encompasses two adjacent columns at a time. Such small filters are unlikely to strongly pick up linkage patterns in the input image data. This is consistent with the results from our ConfuseNN analysis showing that linkage features were not important for the performance of this network.

## CONCLUSION

In summary, we developed ConfuseNN, a method for interpreting the performance of CNNs trained for various population genetic inference tasks. Our method works by shuffling the input haplotype matrix in specific fashions that alter known summary statistics. We applied our method to three published CNNs and found various degrees of sensitivity to feature alterations that: provided insight for improving simulated data design (disc-pg-gan), were consistent with population genetic understanding of the data features known to be important for the certain tasks (Imagene), and underscored the role of architecture features such as filter size (Flagel et al. (2019)). Our method is straightforward to implement and applicable to different algorithms and tasks, providing an accessible and biologically motivated framework for addressing the black-box nature of deep learning algorithms in population genetics.

## METHODS

For each of the three published studies, we replicated the original results using the respective Github repository. For Flagel et al. (2019) and Imagene, we used the provided code to generate simulations for training data and to train the neural networks. We also used the original published procedure to generated simulated test data that underwent shuffling. For disc-pg-gan, we used the trained networks provided by the authors rather than repeating the training procedure. We then used their code to generate simulated test data used for the shuffle tests. We followed the original publications closely, and further details highlighting the specific case we chose from each study are in the Supplementary Methods.

### Software and Data Availability

All code used for analysis in this paper is available at https://github.com/lntran26/ConfuseNN

## Acknowledgments

This work was supported by the National Institute of General Medical Sciences of the National Institutes of Health (R01GM127348 and R35GM149235 to R.N.G.). We thank Sara Mathieson for help with disc-pg-gan and Flagel et al. 2019 and Torada et al. 2019 for making their work easily reproducible.

## Supplementary Methods

### Positive selection classifier CNN - disc-pg-gan

The original study trained several discriminators using different human populations and seed. We analyzed the behavior of a discriminator CNN trained on CEU population data seed 19 from the original publication. This CNN has been fine-tuned with positive selection simulations generated using SLiM.

To generate simulated test data, we used the Riley et al. (2024) SLiM-based procedure to generate 3000 simulated genomic regions for the neutral case (true negative in the binary test) and 2400 simulated genomic regions for positive selection case (true positive), using four positive selection strengths (selection coefficient *s* = 0.01, 0.025, 0.05, 0.1), each contributing 600 simulated genomic regions.

### Positive selection classifier CNN - Imagene

Training and test simulations were generated with *msms* using a three-epoch demographic history model with specified parameters from (Marth et al. 2004) for the CEU population, following Torada et al. (2019). The simulated genomic regions were of size 80 kb and 128 haplotypes were sampled from CEU individuals. The demographic history model has two instantaneous population size change events: a bottleneck of 10, 000 down to 2, 000 individuals at 3500 generations before the present and a recovery and expansion to 20, 000 at 3000 generations before the present. To incorporate selective sweeps, 5000 simulations were generated per selection coefficients *S* = 0, 200, 400 in 2*N* units, which correspond to *S* = 0, 0.01, 0.02.

Each simulation was processed into a 128 × 128 haplotype matrix using major/minor allele polarization and removal of loci with less than 1% allele frequency. The 128 rows in the final matrix correspond to the 128 sampled haplotypes, while the variable number of segregating sites are standardized into 128 with the image resizing python package *scikit-image* (Van der Walt et al. 2014). Both rows and columns of the haplotypes were sorted by frequency of occurrence.

The network used here was implemented using python packages TensorFlow v2.15.0 and Keras v2.15.0. The network architecture consists of three convolutional layers each with 32, 64, and 64 filters respectively with a ReLU activation function followed by a max pooling layer. The output is flattened before being passed into the final output dense layer with 3 nodes corresponding to the 3 classes of selection (0, 200, and 400) and a softmax activation function.

### Demographic history inference CNN

Training and test simulations were generated with *ms* (Hudson 2002) using a three-epoch demographic history model as described in Flagel et al. (2019). Briefly, 100,000 1.5MB regions were simulated with human mutation and recombination rates and demographic parameters randomly sampled from ranges described in Table S1. As done in the original study, we also divided the mutation rate by 10, which is equivalent to randomly downsampling the total number of SNPs by a factor of 10. 80,000 simulations were used in training and 20,000 simulations were used for accuracy testing as well as the scramble test.

Each simulation was processed into a haplotype matrix with 50 rows, corresponding to the number of sampled haplotypes. The width of the matrix equaled the maximum number of segregating sites observed across all simulations. For simulations with fewer sites, columns of 0s were added to the right edges (zero-padding) to make all matrices have the same fixed width.

The position vector has length equal to the maximum number of segregating sites as above minus one. Each value in this vector is the normalized distance between two adjacent segregating sites, with 1 being the total length of the 1.5 MB simulated region. Position vectors are also zero-padded so they all have the same length.

The network used here was implemented using python packages TensorFlow v1.1.0 and Keras v2.0.6. This network has two branches, one takes the haplotype matrix as input and the other takes the position vector as input. The haplotype matrix branch consists of four convolutional layers each with 128 filters followed by a max pooling layer. The position vector branch consists of a dense layer with 32 nodes. The output of these two branches are concatenated before being fed into the final dense layer of 256 nodes. We implemented the best performing hyperparameters reported in the original study, which were 1D convolution type, kernel size 2, output in log scale, drop-out layers throughout, and sorting of chromosomes by genetic similarity.

## Supplementary Results

**Table S1:** Our accuracy scores compare to the original work in Flagel et al. (2019) when replicating the demographic history inference CNN.

|  | Flagel et al. | Our replication | Flagel et al. | Our replication |
| --- | --- | --- | --- | --- |
| Parameter (range) | RMSE | RMSE | Spearman's $\rho$ | Spearman's $\rho$ |
| $N_0(100, 40000)$ | N.A. | 0.584 | 0.59 | 0.591 |
| $N_1(100, 5000)$ | N.A. | 0.764 | 0.54 | 0.544 |
| $N_2(100, 20000)$ | N.A. | 0.571 | 0.84 | 0.856 |
| $T_1(100, 3499.99)$ | N.A. | 0.741 | 0.62 | 0.605 |
| $T_2(T_1 + 1, T_1 + 3500)$ | N.A. | 0.674 | 0.62 | 0.609 |
| RMSE on<br>validation set | 0.43366 | 0.451 |  |  |

**Figure S1:**
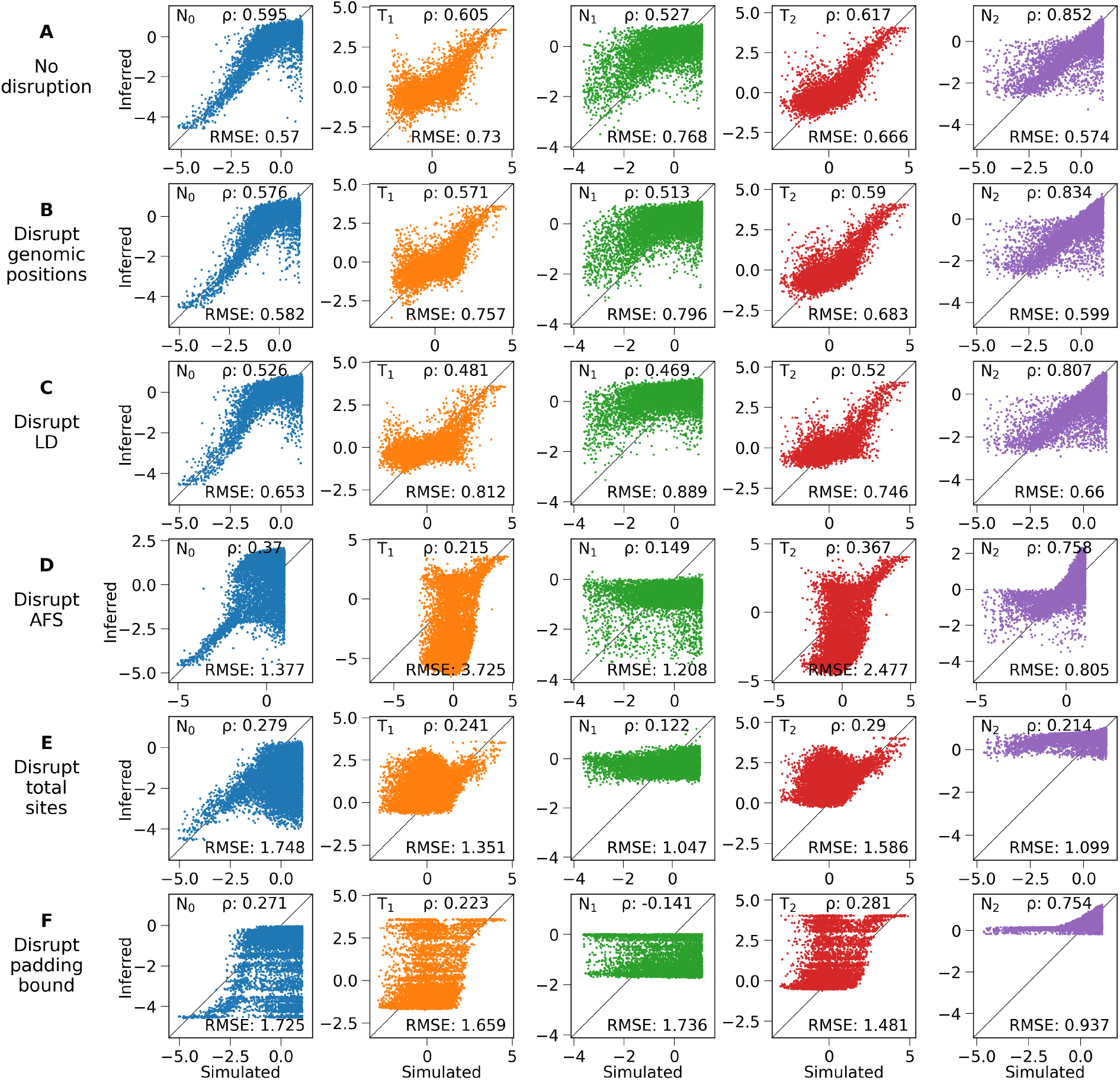
Effects of data shuffling on Flagel et al.’s demographic history inference CNN. (A) Replicating the original CNN’s performance before shuffling. (B-F) The accuracy of the original CNN when tested on different shuffled test data shown in Fig. 1. Accuracy is measured by the Spearman’s rank coefficient *r* and the root mean squared error (RMSE) as per the original work. All parameter values are in log scale.

